# Optimal growth of microbes on mixed carbon sources

**DOI:** 10.1101/120667

**Authors:** Xin Wang, Chao Tang

## Abstract

A classic problem in microbiology is that bacteria display two types of growth behavior when cultured on a mixture of two carbon sources: in certain mixtures the bacteria consume the two carbon sources sequentially (diauxie) and in other mixtures the bacteria consume both sources simultaneously (co-utilization). The search for the molecular mechanism of diauxie led to the discovery of the lac operon and gene regulation in general. However, why microbes would bother to have different strategies of taking up nutrients remained a mystery. Here we show that diauxie versus co-utilization can be understood from the topological features of the metabolic network. A model of optimal allocation of protein resources to achieve maximum growth quantitatively explains why and how the cell makes the choice when facing multiple carbon sources. Our work solves a long-standing puzzle and sheds light on microbes’ optimal growth in different nutrient conditions.

---

During the course of evolution, biological systems have acquired a myriad of strategies to adapt to their environments. A great challenge is to understand the rationale of these strategies on quantitative bases. It has long been discovered that the production of digestive enzymes in a microorganism depends on (adapts to) the composition of the medium (1). More precisely, in the 1940s Jacques Monod observed two distinct strategies in bacteria (*E. coli* and *B. subtilis*) to take up nutrients. He cultured these bacteria on a mixture of two carbon sources, and found that for certain mixtures the bacteria consume both nutrients simultaneously while for other mixtures they consume the two nutrients one after another (2, 3). The latter case resulted a growth curve consisted of two consecutive exponentials, for which he termed this phenomenon “diauxie”. Subsequent studies revealed that the two types of growth behavior, diauxic- and co-utilization of carbon sources are common in microorganisms (4-8). The regulatory mechanism responsible for diauxie, that is the molecular mechanism for the microbes to express only the enzymes for the preferred carbon source even when multiple sources are present, is commonly ascribed to catabolite repression (5, 9-13). In bacteria it is exemplified by the *lac* operon and the cAMP-Crp system (14-17). In yeast, the molecular implementation of catabolite repression differs, but the logic and the outcome are similar (5).

Why have microbes evolved to possess the two strategies and what are the determining factors for them to choose one versus the other? For unicellular organisms, long term survival and growth at the population level are paramount. In the exponential growth phase, cell optimizes growth by optimally allocating its resources (18-24). In particular, Hwa and colleagues developed a model of optimal growth with constraints on protein resource (20, 21). In this paper, we extend this approach to address the question of multiple carbon sources and show that the two growth strategies can be understood from optimal growth further constrained by the topological features of the metabolic network.

## Categorization of Carbon Sources

Carbon sources taken by the cell serve as substrates of the metabolic network, in which they are broken down to supply pools of amino acids and other components that make up a cell. Note that amino acids take up a majority of carbon supply (about 55%) (25-27). As shown in Fig. 1, different carbon sources enter the metabolic network at different points (27). Denote those sources entering the upper part of the glycolysis Group A and those joining at other points of the metabolic network Group B (Fig. 1). Studies have shown that when mixing a carbon source of Group A with that of Group B, the bacteria tend to co-utilize both sources and the growth rate is higher than that with each individual source (6, 7, 28). When mixing two sources both from Group A, the bacteria usually utilize a preferred source (of higher growth rate) first (4, 6, 11, 13, 29, 30).

**Fig. 1.**
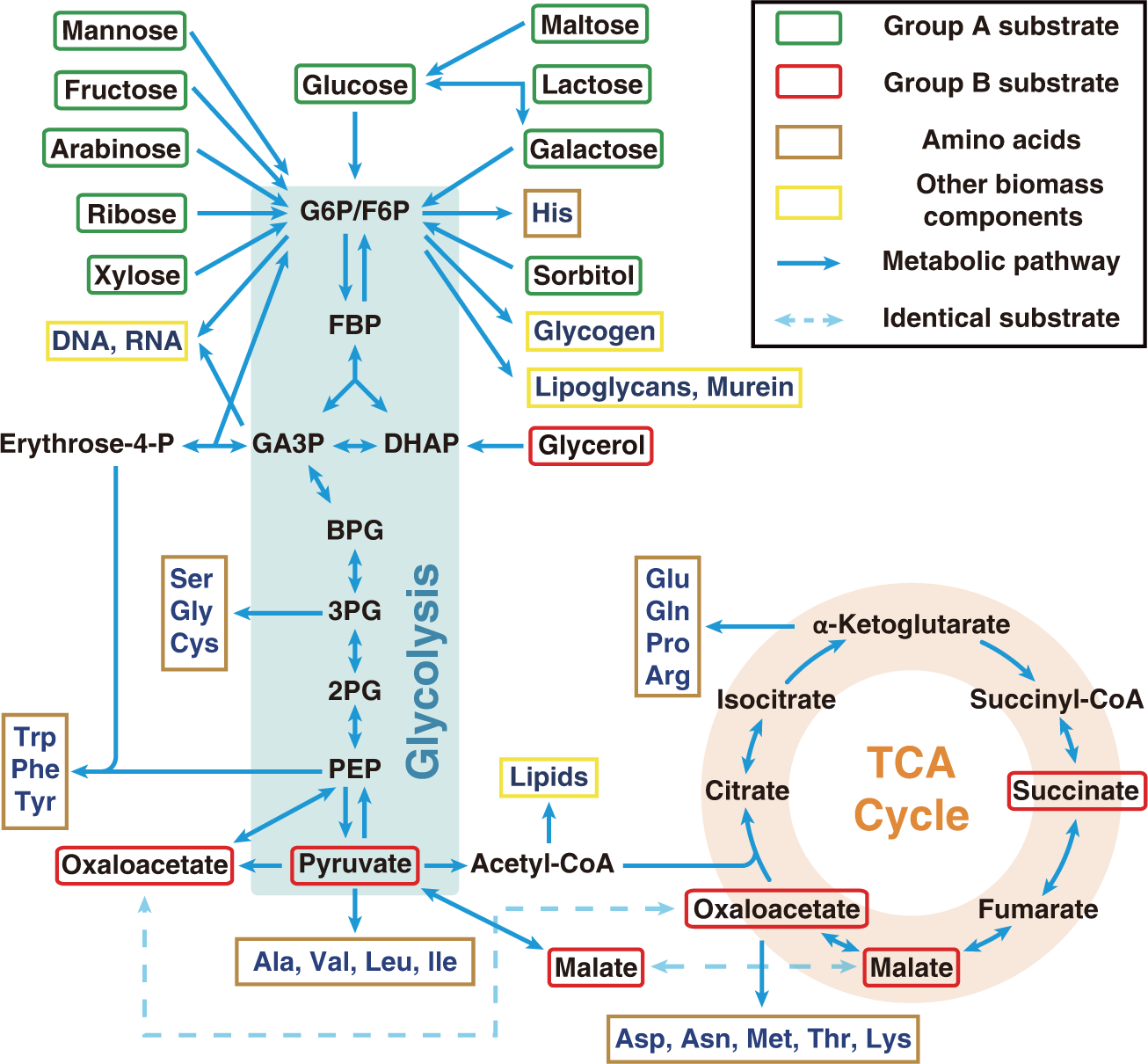
Metabolic network of carbon source utilization. Group A substrates (in green squares) can be simultaneously utilized with Group B substrates (red squares), whereas substrates paired from Group A display diauxie. Only major pathways are shown here.

## Origin of Diauxie for Carbon Sources in Group A

Let us first consider the case in which both carbon sources are from Group A. In this case, if we group the precursors of biomass components (amino acids and others) into various pools, then all these pools lie downstream of the carbon sources (Fig. 1). The topology of the metabolic network is then equivalent to Fig. 2A (see Methods), where *A1* and *A2* can be any two carbon sources from Group A. We proceed to solve this simple model using an optimization principle (20, 21) (see Methods).

**Fig. 2.**
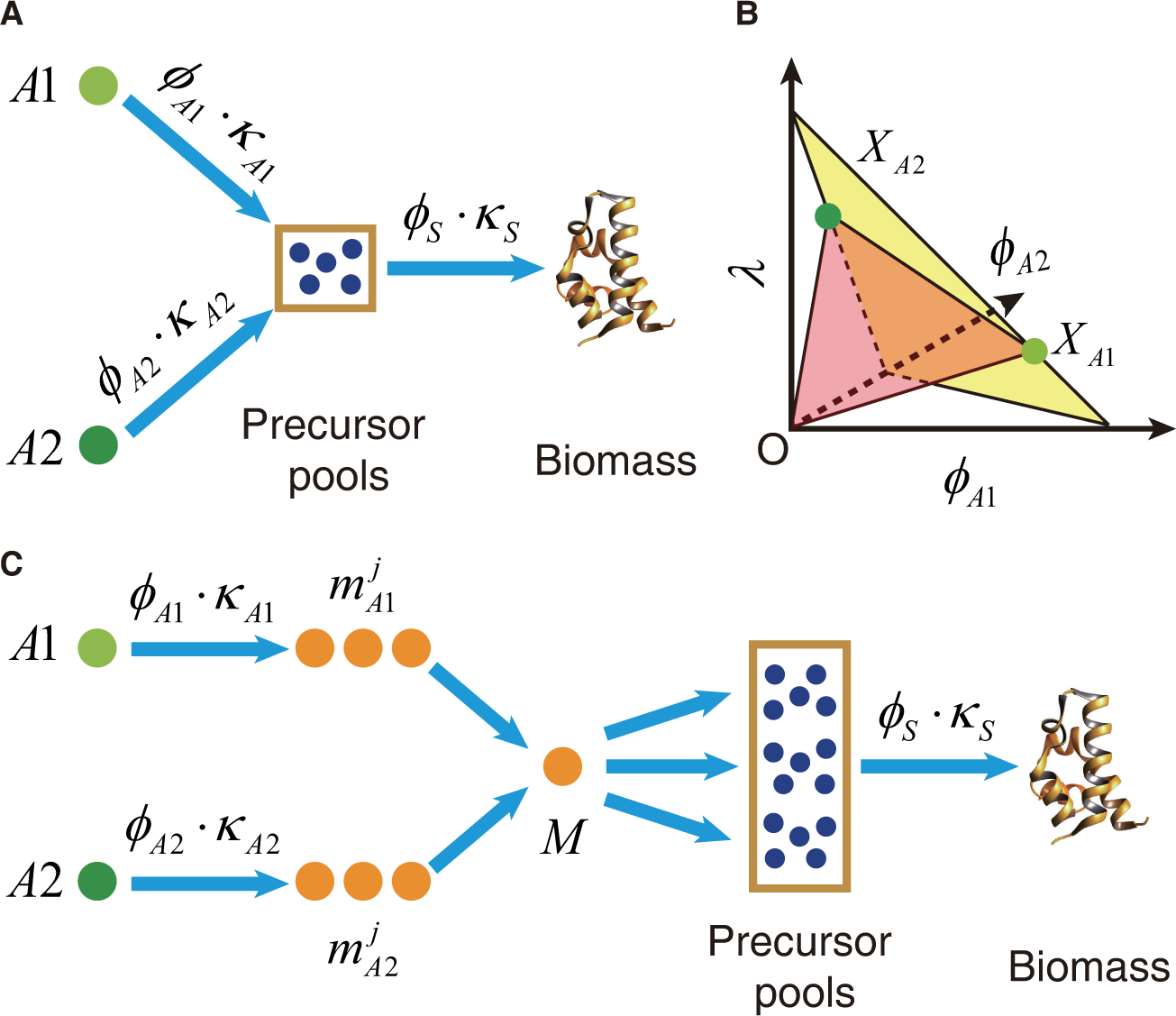
The origin of diauxic growth. (A) Minimal model of diauxie. The carbon sources *A*1 or *A*2 or both can supply the precursor pools. The cell grows faster if only the more efficient source is utilized as shown in (B). (B) The relations among the enzyme mass fractions and growth rate. The maximal growth rate is at the apex (green points) *X_A1_* (*ϕ_A2_*=0) or *X_A2_* (*ϕ_A1_*=0). In either case, the suboptimal substrate is not consumed. (C) Topology of metabolic network with two Group A sources. The two carbon flux pathways from sources *A*1 and *A*2 can have multiple intermediate nodes (metabolites)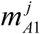 and 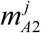 before merging to a common node *M*, after which the flux is diverted to various precursor pools.

In Fig. 2A, enzymes carrying and digesting nutrient *Ai* (*i*=1, 2) into the precursor pools are simplified to a single enzyme *E_Ai_* with protein mass fraction 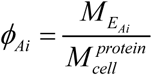, where *M_E_Ai__* is the total mass of the enzyme *E_Ai_* and 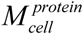 the total protein mass in the cell. The carbon flux to the precursor pools from source *Ai* is proportional to (*ϕ_Ai_* and takes the Michaelis-Menten form (see Supporting Information for details): *J_Ai_* = *ϕ_Ai_*. *k_Ai_*, where 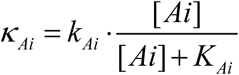 (which we denote as the nutrient quality) with [*Ai*] being the concentration of *Ai*. We define nutrient efficiency of a carbon source as the carbon flux delivered divided by the protein mass faction dedicated to the delivery. In this model, it is simply *J_Ai_*/*ϕ_Ai_* = *K_Ai_*. Pools of precursors are utilized to manufacture biomass with a flux rate proportional to the growth rate of the cell: *λ* = *ϕ_s_* · *K_s_*, where *ϕ_S_* represents the protein mass fraction of the enzymes dedicated in synthesizing biomass from the precursor pools, and *K_S_* a kinetic constant (see Supporting Information for details). Since the ribosomes by far constitute the majority of the enzymes for biomass synthesis, *ϕ_S_* is dominated by *ϕ_R_*, the mass fraction of the ribosomes and *K_s_* ≈ *K_t_*, where *K_t_* is a parameter determined by protein translation rate (20, 21). The protein mass and carbon flux constraints give (see Supporting Information for details)

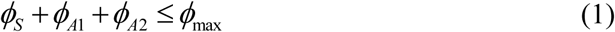

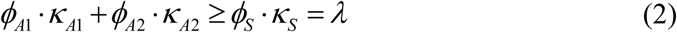

Where *ϕ*_max_ is a positive constant less than 1, representing the largest proportion of protein mass that can be allocated to *ϕ_S_, ϕ_A1_* and *ϕ_A2_*. These relations are depicted in a 3-dimensional graph (Fig. 2B), with *ϕ_A1_* the x-axis, *ϕ_A2_* the y-axis and *λ* (=*ϕ_s_*.*K_S_*) the z-axis. The yellow plane corresponds to the upper bound of Eq. 1, while the light red plane the upper bound of Eq. 2. *ϕ_A1_, ϕ_A2_* and *λ* should all be non-negative. Under these conditions, the optimal solution (solution with maximal *λ*) should be at either *X_A1_* or *X_A2_* (Fig. 2B). If *K_A1_* > *K_A2_*, then *X_A1_* is the optimal point with 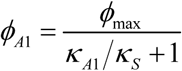 and *ϕ_A2_* = 0 (see Supporting Information for details), which means that the cell expresses only the enzyme for A1 and thus utilizes only A1. Conversely, if *K_A1_* < *K_A2_*, the optimal solution is *X_A2_* and the cell utilizes only A2. In either case, cells only consume the preferable carbon source, which corresponds to the case of diauxie (2, 3, 6, 9, 11).

In the above coarse-grained model, the nutrient efficiency of the carbon source *Ai* is lump summed in a single effective parameter *K_Ai_*. In practice, there are intermediate nodes and enzymes along the pathway as depicted in Fig. 2C. So in order to make comparison between the actual carbon sources, one should take into account the cost of enzymes for the intermediates. Denote 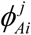 the enzymes catalyzing the intermediate nodes 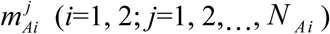, and define 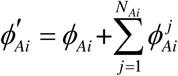, which is the total fractional protein mass for enzymes dedicated to the branch *Ai*→*M*. The nutrient efficiency for the branch is then 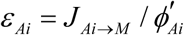, where *J_Ai→M_* is the carbon flux from *Ai* to *M*(*i*=1,2). Assuming that the flux is conserved along the branch so that 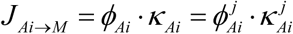, (*j*=1, 2,…, *N_Ai_*), where 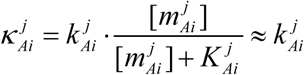 is the substrate quality of 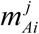, we have (see Supporting Information for details)

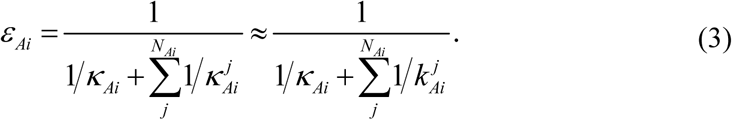

The equivalent of Eqs. 1 and 2 is

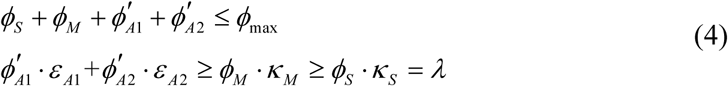

For *ε_A1_* > *ε_A2_*, the optimal solution is *ϕ_A1_* = *λ/K_A1_*, 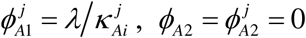*ϕ_M_* =*λ/k_M_*, and 
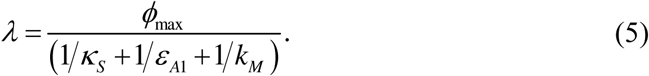

Only the nutrient (*A*1) with higher efficiency (*ε*_*A*1_) is utilized. Note that the growth rate is the same as when cultured with *A*1 alone.

## Ratio Sensing

We have shown that when there are two Group A sources *A*1 and *A*2 available, the cell will only utilize the one with higher efficiency. However, note that the nutrient efficiency *ε_Ai_* (Eq. 3) depends on the concentration of the nutrient [*Ai*] through the nutrient quality 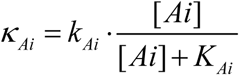. *ε_Ai_*([*Ai*]) decreases with *[Ai]*. Thus an initially preferred carbon source can become less preferable when its concentration becomes too low. This is because for lower nutrient concentration more enzymes have to be used to deliver the same carbon flux. Suppose that *A*1 is a preferred carbon source than *A*2 when both are abundant (*ε_A1_ > ε_A2_*, for [*Ai*] > *K_Ai_* (*i* = *1,2*)). For a fixed [*A2*], when the concentration of *A1* drops below a value [*A1*]_T_such that *ε_Al_([A1]_T_) = ε_A2_([A2]), A*2 becomes more efficient and should be utilized instead of *A*1. From Eq. 3, the turning point is given by (see Supporting Information for details) 
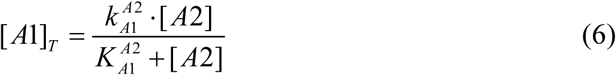
where 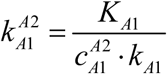 and 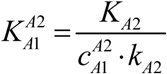, in which the parameter 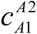 is defined as 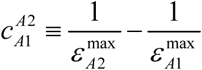, with 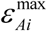 the maximum branch efficiency of the nutrient *Ai* (at saturating nutrient concentration). For 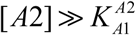, the turning point 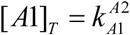 does not depend on [*A*2]. For 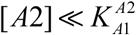, the turning point is reduced to 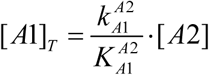, a form of ratio sensing. That is, the cell will sense not the absolute concentration of [*A1*] and [*A2*], but their ratio, to make the decision. Ratio sensing was recently observed in the budding yeast *Saccharomyces cerevisiae* cultured in glucose-galactose mixed medium (29). The measured turning point is in quantitative agreement with Eq. 6 (Fig. 3).

**Fig. 3.**
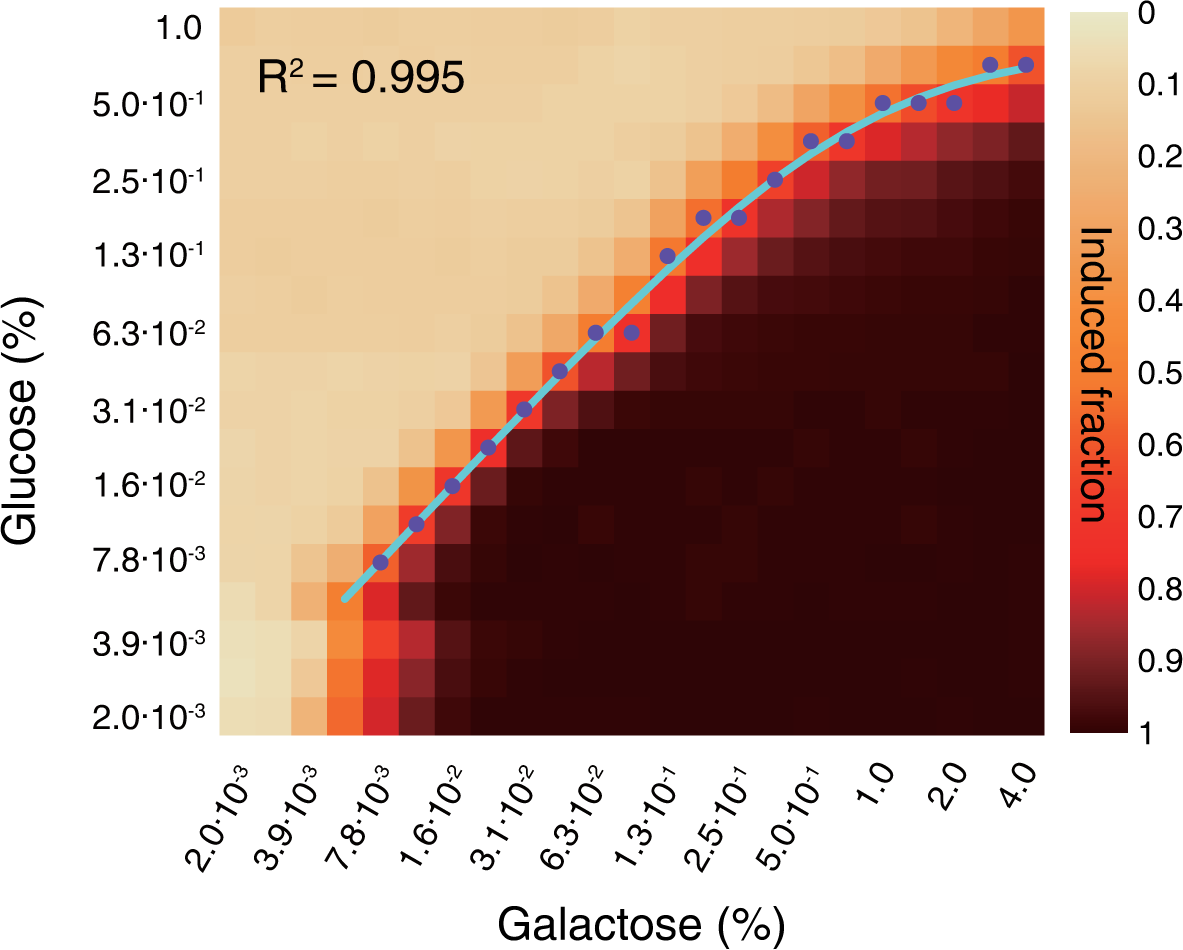
Concentration dependence of turning point. In the experiment of Escalante-Chong et al. (29), yeast cells were cultured with a mixture of glucose and galactose of various combinations of concentrations. The induction of galactose pathway was measured in single cells with flow cytometry. The heat map represents the fraction of cells with the galactose pathway turned on for given pairs of concentrations (Reproduced with permission). The purple dots indicate the glucose concentration at which the induction fraction is at or just above 0.5 for given galactose concentration. The solid line is a fit with Eq. 6 ( *R^2^* = 0.995,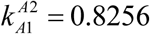 and 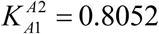).

## Co-Utilization of Carbon Sources

The diauxic growth is due to the topology of the metabolic network, in which Group A sources enter the network in the upper part of the glycolysis and converge to a common node (G6P/F6P) before diverting to various precursor pools (Figs. 1 and 2C). The situation is different if the two mixed carbon sources are from Groups A and B, respectively (denoted as “A+B”). (Some combinations of two Group B sources also fall into this category and can be analyzed similarly; see Fig. S2C.) Group B sources can directly supply some precursor pools without going through the common node (G6P/F6P) (Fig. 1). As an example, the topology of the metabolic network in the presence of one Group A source in combination with the Group B source succinate is shown in Fig. 4A. More examples of “A+B” are shown in Fig. S2. All “A+B” cases can be mapped to a common coarse-grained model depicted in Fig. 4B, although the actual position of nodes *M* and *N* in the metabolic network, and the contents of Pools 1 and 2 may depend on each specific case. As obvious from Fig. 4B, source *A* or *B* alone could in principle supply all precursor pools. However, because of the location of the precursor pools relative to the sources, it may be more economical for one pool to draw carbon flux from one source and the other from the other source.

**Fig. 4.**
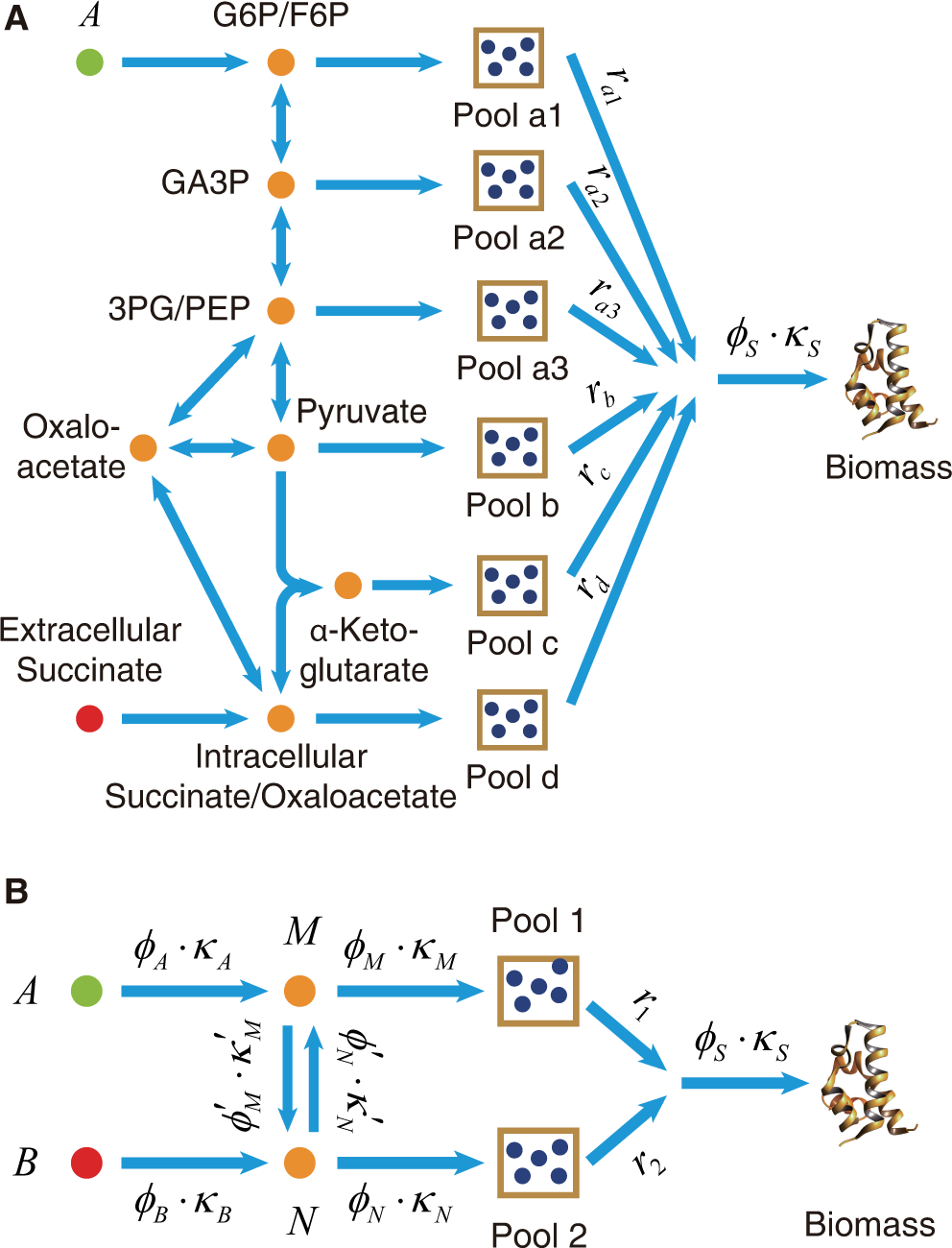
Network topology behind co-utilization. (A) Topology of the metabolic network when Group A source is mixed with a Group B source succinate. (B) Minimal model of co-utilization. In synthesizing biomass, the two precursor pools supply *r_1_* and *r_2_* carbon flux, respectively. Either pool can draw flux from either of the two sources *A* and *B*. Under certain conditions, it is optimal for different sources to supply different pools exclusively.

In the model, the two pools of precursors supply *r*_1_ and *r*_2_ carbon flux respectively to the synthesis of biomass. Two intermediate nodes *M* and *N* can interconvert to each other with the respective enzymes 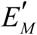(of protein mass fraction 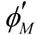) and 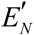(of protein mass fraction 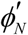). To determine which of the two carbon sources should supply which pool(s), we apply branch nutrient efficiency analysis. For Pool 1, we compare the efficiency of *A* and *B* in supplying flux to node *M*; while for Pool 2 to node *N*. The results are

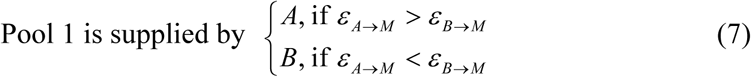

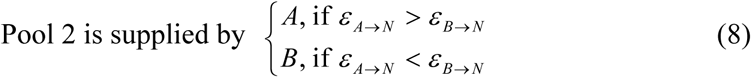
where 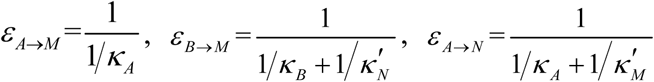 and 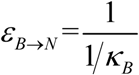.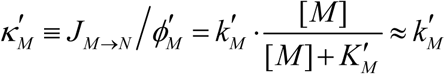 and 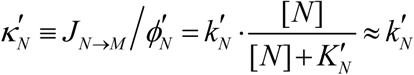 are the substrate quality of *M* and *N* in reactions with enzyme 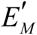 and 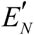, respectively (see Supporting Information for details). It is easy to see from inequalities 7 and 8 that if the following condition is met 
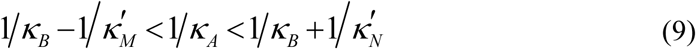
then *A* supplies Pool 1 and *B* supplies Pool 2 ‐‐ the two carbon sources are simultaneously consumed. In reality, there are multiple intermediate nodes in between the *M-N* interconversion (Figs. 1, 4A and S2). Similar to Eq. 3 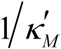 and 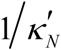 here actually represent summations of all intermediate terms between *M* and *N* in the metabolic network.

If *A* and *B* are co-utilized, the growth rate is (see Supporting Information for details) 
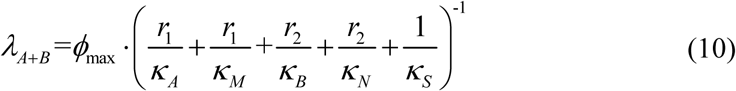

In comparison, with a single carbon source (*e. g. A*), the growth rate is 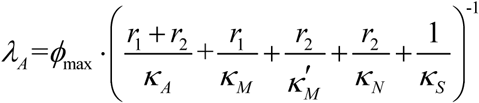 As 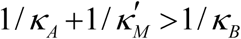 (see Eq. 9, the condition for co-utilization), *λ_A+B_ > λ_A_*.

## Carbon Source of Biomass components

In order to apply the above analysis to the real case, we collected the available data for metabolic enzymes from the literature (Table S1), and calculated the values of the branch nutrient efficiency from several carbon sources to the metabolites F6P, GA3P, 3PG, PEP, pyruvate or oxaloacetate, which correspond to the nodes *M* or *N* in the simplified network of Fig. 4B (see also Figs. 1, 4A and S2). The results are shown in Table S2. Applying the analysis of Eqs. 7 and 8 to the real network, one can identify the carbon source supplier of each amino acid pool and other precursors pools under optimal growth (Table S3).

## Discussion

The topology of the metabolic network can be simplified to coarse-grained models shown in Figs. 2A and 4B for the analyses of diauxie and co-utilization of carbon sources. The sources of Group A all go through a common intermediate node. Thus they compete for delivering carbon flux to the precursor pools. The more efficient one wins (22). It has been observed that there is a hierarchy among Group A sources ranked according to the growth rate on single carbon source – when two or more sources are present the bacteria use the one that delivers the highest growth rate (6, 30). This is a natural consequence of our theory. As can be seen from Eq. 5, a higher growth rate implies a higher nutrient efficiency and thus a higher priority to be utilized. How this hierarchy is implemented molecularly is a very interesting question (13). Ratio sensing is another consequence of our theory. It remains to be seen experimentally whether it is widely implemented for all pairs of Group A nutrients and across microbes. It could well be that the microbe cares only about the most frequently encountered (or the most important) combinations of nutrients and would not invest resources to ratio sense the others.

When Group B source is present along with Group A source, it can take a shortcut to reach some of the precursor pools (Fig. 4) and can be more efficient to supply these pools. (Some combinations of two Group B sources may face the same situation and thus can be co-utilized.) An experimental test for our theory of co-utilization is to verify/falsify Table S3. There is no data yet in the literature to compare directly with Table S3. There are, however, experimental data on the relative fractions of fluxes the cells are drawing from the two carbon sources when cultured on sources A and B (6, 30). These quantities can be estimated with the knowledge of Table S3 and the composition of amino acids in a cell (27). The results are consistent with experimental data (Fig. 5). In a recent experiment with *Methylobacterium extorquens* AM1 (28), the carbon source of certain metabolites were traced with isotope labeling in a co-utilization case. The outcome is consistent with our theory (see Supporting Information for details).

**Fig. 5.**
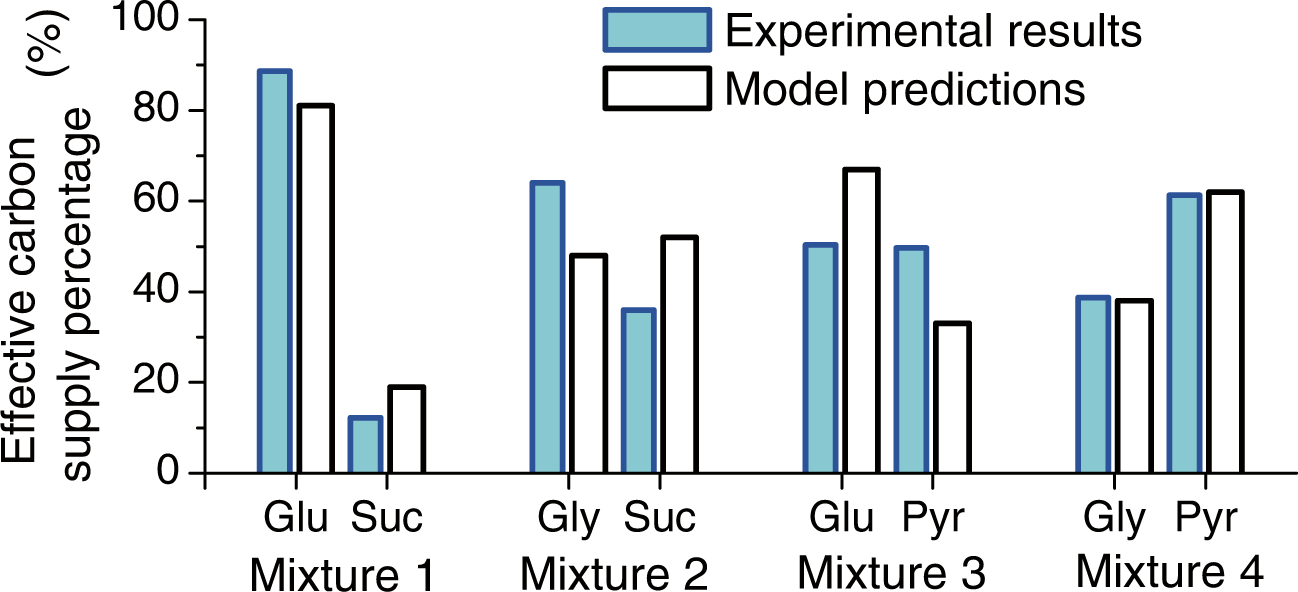
Carbon source supply percentages in the cases of co-utilization. The experimental results were estimated from published data (6), while the model prediction values were calculated from the composition of biomass in a cell and the predicted source suppliers (Table S3) of the biomass components.

Cell growth is a fundamental issue in biology. The present work deals with relatively stable growth conditions and the simple exponential growth behavior. In this case, there is a body of experimental evidence for optimal growth (maximum growth rate) (18-24). In reality the environment the microbes face can be highly variable and uncertain. Their long-term “fitness” of the population may not simply be determined only by the growth rate of individual cells in the exponential phase, but a result of trade-offs that best adapt to the changing environment. Strategies such as bet hedging, memory of the past and anticipation of the future are found to exist in microorganisms (31-40). Furthermore, while the phenomena of diauxie versus co-utilization are widely spread in microbes, they are bound to be variations and exceptions. The nutrient uptake strategy or eating habit of a microbe is shaped by its environmental history. For example, certain microbes may have different hierarchies of preferable carbon sources (4). One challenge is to understand how cells and population behave in and evolve with the environment in a general and quantitative framework.

Another interesting point is cell-to-cell variability. What our theory gives is the average behavior. However, the behavior of individual cells can be variable. For example, in the ratio sensing experiment we discussed before (29), the turning point to switch on the galactose pathway is variable from cell to cell, and in each cell the switching is an all-or-none transition (bistable with memory). The bistability and perhaps at least some of the variability in the switching point may well be the outcome of evolution to cope with environmental fluctuations and uncertainties. Interestingly, our prediction of the turning point agrees very well with the average behavior of the population (Fig. 3). A challenge in future research would be to quantitatively understand the variance.

## Methods

Coarse graining of the metabolic network is done in such a way as to preserve the network topology but grouping metabolites, enzymes and pathways into single representative nodes and corresponding effective enzymes. In particular, a linear pathway is lump summed into two nodes (start and end) connecting with a single effective enzyme.

The protein resource allocation model is based on the work of Hwa and colleagues (20, 21). For our purpose here, the proteins in a cell are partitioned into three classes: carbon catabolic enzymes (C), biomass synthesizing enzymes (S) and everything else (Q). The masses of the three classes add up to the total protein mass in a cell: 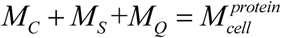, or *ϕ_C_* + *ϕ_S_* + *ϕ_Q_* = 1, where 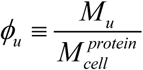 (*u* = *C, S, Q*) is the protein mass fraction. Optimal growth is achieved by optimally distribute *ϕ_C_* and *ϕ_S_* under the constraint *ϕ_C_* + *ϕ_S_* = 1 - *ϕ_Q_* ≤ *ϕ_max_*. With multiple carbon sources, *ϕ_C_* is broken down to subgroups according to the sources and pathway topologies as shown in the main text.

## Author Contributions

X. W. and C. T. designed the study, developed the model, and wrote the paper. X. W. carried out the analysis.

## Acknowledgements

We are grateful to Yuan Yuan and Haoyuan Sun for their insights. We thank Xiaojing Yang, Shanshan Qin, Yimiao Qu and Chang Chang for helpful discussions. This work was supported by Chinese Ministry of Science and Technology (2015CB910300) and National Natural Science Foundation of China (91430217).

